# Decoding *Anadara* shell morphology with deep learning

**DOI:** 10.64898/2026.04.17.719170

**Authors:** Masato Tsutsumi, Nen Saito, Tomohiro Yamaguchi, Takenori Sasaki, Chikara Furusawa

## Abstract

Accurate shell shape quantification is critical for studying biodiversity and evolution, yet intraspecific variability in bivalves makes morphology-based identification difficult. Traditional methods, including landmark-based analyses and elliptic Fourier descriptors, suffer either from subjectivity in homologous point selection or from limited use of contour information. Here, we introduce Morpho-VAE, a deep generative framework integrating a variational autoencoder with a supervised classifier, to analyze shell images of five *Anadara* species. Morpho-VAE outperforms conventional approaches in species classification by embedding morphological variation into a low-dimensional space where species cluster distinctly. To highlight species-specific morphological patterns, we develop a patch masking assay, revealing the hinge line as a shared morphological marker across species and species-specific regions near the umbo and anterior ventral margin. The decoder further enables morphological visualization via image reconstruction and interpolation. Our results show that Morpho-VAE can automatically extract species-defining morphological patterns from raw images, providing complementary or novel insights beyond traditional morphometric methods.

## INTRODUCTION

The diversification of shell morphology has long been studied in taxonomy and ecology because shells are easily preserved as fossils, allowing large-scale sampling of both extant and fossil species [1]. Bivalves lead sessile or relatively immobile benthic lifestyles, meaning the physicochemical conditions of their habitat directly influence shell growth [2]. Consequently, even within the same species, significant morphological variations are observed because of differences in habitat substrate [3–5], latitude [6, 7], and other factors [8, 9]. However, such morphological variation also reduces the robustness of species identification and classification based on morphology [10, 11]. While a quantitative parameterization framework using the characteristics of coiling has been established for gastropod shells [12–15], this framework cannot be applied to bivalve shells, which rarely have coiling characteristics. Parameterization with a low-dimensional parameter for bivalve shells is not straightforward [16]; however, it is important for understanding which morphological characteristics capture shell shape dispersion and for conducting an interpretable analysis of bivalve diversification and evolution in parameter space.

Geometric morphometrics has been used to quantify bivalve shapes. The landmark method has been extensively applied to analyze bivalve morphologies [3–9, 17–19]. In this method, morphological diversification depending on environmental factors is projected onto the low-dimensional parameter space obtained through dimension reduction. This space is known as morphospace. Based on this morphospace, shape distribution has been examined at both the intraspecific [3, 4, 6, 7] and interspecific levels [5, 9, 17, 18]. However, landmark methods require researchers to subjectively set landmark positions and determine their number, which affects whether the shape variation can be adequately characterized [20]. Furthermore, it is difficult to compare species for which homologous points are not clearly defined. Edie et al. noted that bivalve shells are a classic example of homologous structures with few universally homologous points [16]. Elliptic Fourier analysis (EFA) is a widely used landmark-free method. This analysis, based on extracted shell contours, has been used to investigate the morphologies of both extant species [21–25] and fossils [26–29]. However, EFA is restricted to analyzing contour information and tends to average local variations and fine structures [30]. Thus, introducing novel measurement techniques distinct from existing methods suggests the potential to extract morphological features that are difficult to capture using conventional approaches.

In recent years, deep learning methods, specifically convolutional neural networks (CNN), have become a new trend in morphological analysis, including the quantification of inter- and intraspecific variation [31–35]. This approach uses image data as input, eliminating the need for manual landmark definition and enabling objective and comprehensive information acquisition. To analyze bivalve morphology, Hofmann et al. explored supervised multi-level classification and unsupervised similarity learning (SimCLR) to infer taxonomic affinities and genetic distances from bivalve images [31]. This study focused on classification tasks; therefore, the visualization and interpretation of morphospace were not investigated. Other than bivalve research, Cuthill et al. used a deep convolutional triplet network (ButterflyNet) to quantify the phenotypic similarity and dissimilarity of butterflies through 64-dimensional spatial embeddings [33], enabling classification of *Heliconius* species. While this study constructed a latent space reflecting phylogenetic relationships and visualized it in a low-dimensional space, a systematic examination of how morphological variation is organized across this space was beyond the scope of that work. Our previously proposed Morpho-VAE integrates a variational autoencoder (VAE) [36] with a classifier. This approach compresses high-dimensional data into a low-dimensional latent space while preserving the original image information, enabling the automatic extraction of morphological features [37]. Moreover, the generative ability of the model via the decoder structure enables the reconstruction of an image from any point in this low-dimensional latent space. This allows the visualization of continuous morphological changes along the latent space, thus significantly enhancing the interpretability of the space. This approach has been used to analyze mandible shapes [37] but has not been applied to bivalve shapes.

In this study, we applied Morpho-VAE to classify five species of *Anadara*, specifically focusing on identifying diagnostic morphological features from raw images. Although Morpho-VAE constructs a latent space representing global shape variations, identifying specific local regions that contribute to species discrimination remains challenging. To address this, we implemented a quantitative patch-based analysis to evaluate the contribution of local morphological regions to classification. By systematically masking specific parts of the shell image and monitoring the changes in classification performance (e.g., F1 score), we pin-pointed the local areas that are essential for distinguishing the species. We compared these deep learning-derived features with those obtained using landmark methods and EFA. Using this approach, we demonstrated that our method can detect subtle and localized morphological determinants that are often overlooked or difficult to quantify using conventional morphometrics.

## RESULTS

Here, we applied Morpho-VAE to the image data of five species of the genus *Anadara*: *Anadara broughtonii* (*N* = 20), *Anadara inaequivalvis* (*N* = 127), *Anadara satowi* (*N* = 22), *Anadara kagoshimensis* (*N* = 35), and *Anadara ferruginea* (*N* = 26) (Fig. 1a). The procedures for image acquisition and preprocessing are shown in SFig. 5 and SFig. 6. The architecture of Morpho-VAE [37] comprises a combination of VAE structures, including the encoder and decoder with a classification module connected to the latent space (SFig. 8). This combined structure provides a nonlinear supervised dimension reduction while preserving the ability of image reconstruction. The networks were trained using 256 *×* 256-pixel grayscale image data with appropriate image alignment (see Methods). The dataset was divided into the training/validation/test data in a ratio of 3:1:2. This overall allocation was obtained by a group-aware split that first assigned approximately 67% vs. 33% of specimens within each class to training versus test, then subdivided the training portion into approximately 75% vs. 25% for training versus validation (Methods), yielding the 3:1:2 ratio across all specimens. Hereafter, only models with training accuracy above 90% were analyzed. The learning curves are shown in SFig. 9, indicating successful training processes.

**Fig. 1.**
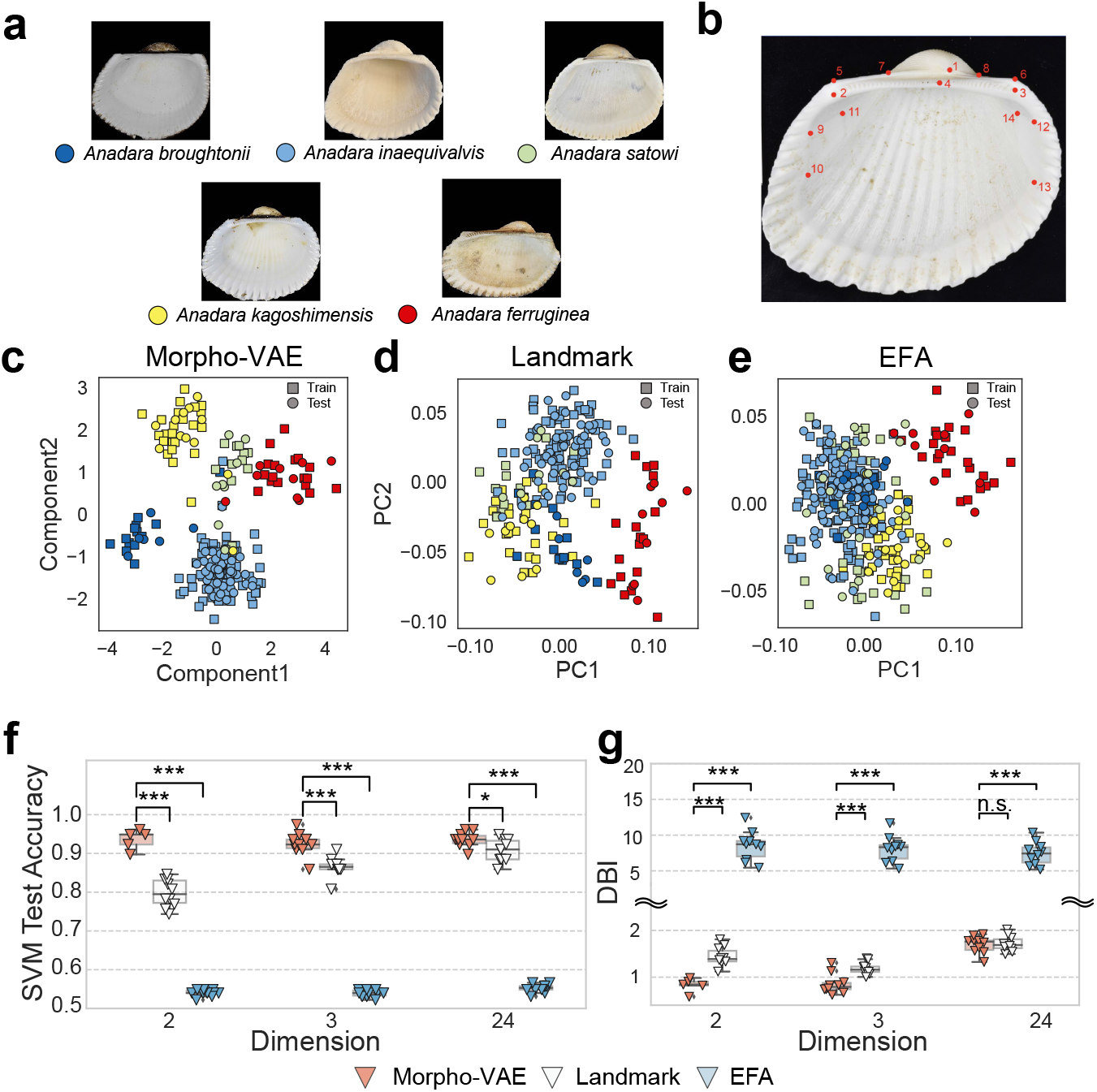
**a**, Representative shell images of the five *Anadara* species. **b**, Left valve of *Anadara inaequivalvis* with landmarks from 1 to 14. The locations of the marked landmarks are as follows: 1. Umbo, 2. Posterior end of the hinge line, 3. Anterior end of hinge line, 4. Commissure of anterior and posterior hinge tooth, 5. Posterior dorsal corner, 6. Anterior dorsal corner, 7. Posterior end of the beak, 8. Anterior end of the beak, 9. Dorsal junction of posterior muscle scar and pallial line, 10. Ventral junction of posterior muscle scar and pallial line, 11. Dorsal end of posterior muscle scar, 12. Dorsal junction of anterior muscle scar and pallial line, 13. Ventral junction of anterior muscle scar and pallial line, 14. Dorsal end of anterior muscle scar. **c-e**, Two-dimensional latent space distributions from Morpho-VAE (**c**), landmark-based PCA (**d**), and EFA-based PCA (**e**). The squares and circles represent the training and test data, respectively. Each color corresponds to a different species. **f-g**, Quantitative comparison of separability in latent space across different methods. **f**, SVM test accuracy for species classification. Each data point corresponds to the result with one of the ten different train/test data combinations. **g**, Davies–Bouldin index (DBI) as a clustering separation index, where values smaller than one indicate well-separated clusters. Statistical significance is denoted as *:*p* < 0.05; **: *p* < 0.01; ***:*p* < 0.001; n.s.: *p ≥* 0.05.

For visualization in Fig. 1c, we constructed a two-dimensional latent space using Morpho-VAE. The Morpho-VAE latent space showed clear clustering across all species for both the training (squares in Fig. 1c) and test datasets (circles). For comparison, we applied a landmark-based approach, where the *x* and *y* coordinates of 14 landmarks (Fig. 1b) were reduced to two dimensions using principal component analysis (PCA). To ensure consistency with Morpho-VAE, PCA was performed only on the training data. Fig. 1d illustrates the latent space constructed by PCA: the squares denote the training data, and the circles denote the test data projected onto the same plane (see Methods). In this landmark-based latent space, *A. ferruginea* (red) and *A. inaequivalvis* (blue) were separated, but the remaining species showed substantial overlap. We further conducted EFA, in which the shell outline was approximated using up to the 20th Fourier mode, followed by dimensionality reduction with PCA (see Methods). In the EFA latent space, *A. ferruginea* again exhibited a distinct separation from other species, but the remaining species displayed partial overlap and poorly defined boundaries, highlighting the limitations of this method for species identification.

By quantitatively evaluating the species-separation performance in the latent space of each method, we demonstrated that Morpho-VAE outperformed the other two methods. First, we evaluated the degree of cluster separation using a support vector machine (SVM). SVM solves classification tasks and can provide a high classification accuracy for data points showing a clear separation of differently labeled data in the latent space, indicating that the classification accuracy can be used as a measure of cluster separation. Latent space construction and SVM training were performed based on the training data. Fig. 1f shows the SVM test accuracy, where each data point corresponds to the result of one of the 10 train/test data combinations (see Methods). Morpho-VAE showed significantly higher accuracy than that of the other methods for the prescribed latent dimensions *N*_*z*_ = 2, 3, and 24 (*p*< 0.05) (Fig. 1f), indicating higher species-separation performance. In addition, we calculated a clustering separation index, the Davies–Bouldin index (DBI) [38], which takes lower values, indicating better-separated clusters (see Methods). Morpho-VAE showed DBI < 1 for *N*_*z*_ = 2 and 3, whereas Landmark and EFA exhibited DBI > 1 for all examined dimensions (Fig. 1g). These results show that Morpho-VAE can embed image data into a low-dimensional latent space while preserving local morphological information, demonstrating a clear advantage in image-based automatic analysis without defining landmark configurations.

### Morphological features of each method

Based on the results shown in Fig. 1, we visualized how the landmark method, EFA, and Morpho-VAE captured morphological features. In the landmark method, for each landmark position (from 1st to 14th), we calculated the average coordinates over all samples and the average over samples of each class and defined displacement vectors between them. For instance, large displacements were observed for *A. ferruginea* at landmarks 5 and 11 (red arrows in Fig. 2a at landmarks 5 and 11), which characterizes the morphology of that species. This is consistent with the results in Fig. 1d, where *A. ferruginea* has a distinctly higher PC1 value, and PC1 has large components in the 5th, 7th, and 11th landmarks (Fig. 2b). The directions of the displacement vectors at landmarks 5 and 6 for *A. ferruginea* (red arrows in Fig. 2a at landmarks 5 and 6) can be explained because *A. ferruginea* exhibits a remarkable elongation of the dorsal corner (a line along landmarks 5 and 6) compared to other species (Fig. 1a). *A. kagoshimensis* has a distinctly negative PC1 value (Fig. 1d) compared with that of *A. ferruginea*. This is reflected by the large displacement vector for *A. kagoshimensis* at landmark 5 (Fig. 2a; yellow arrow at landmark 5).

**Fig. 2.**
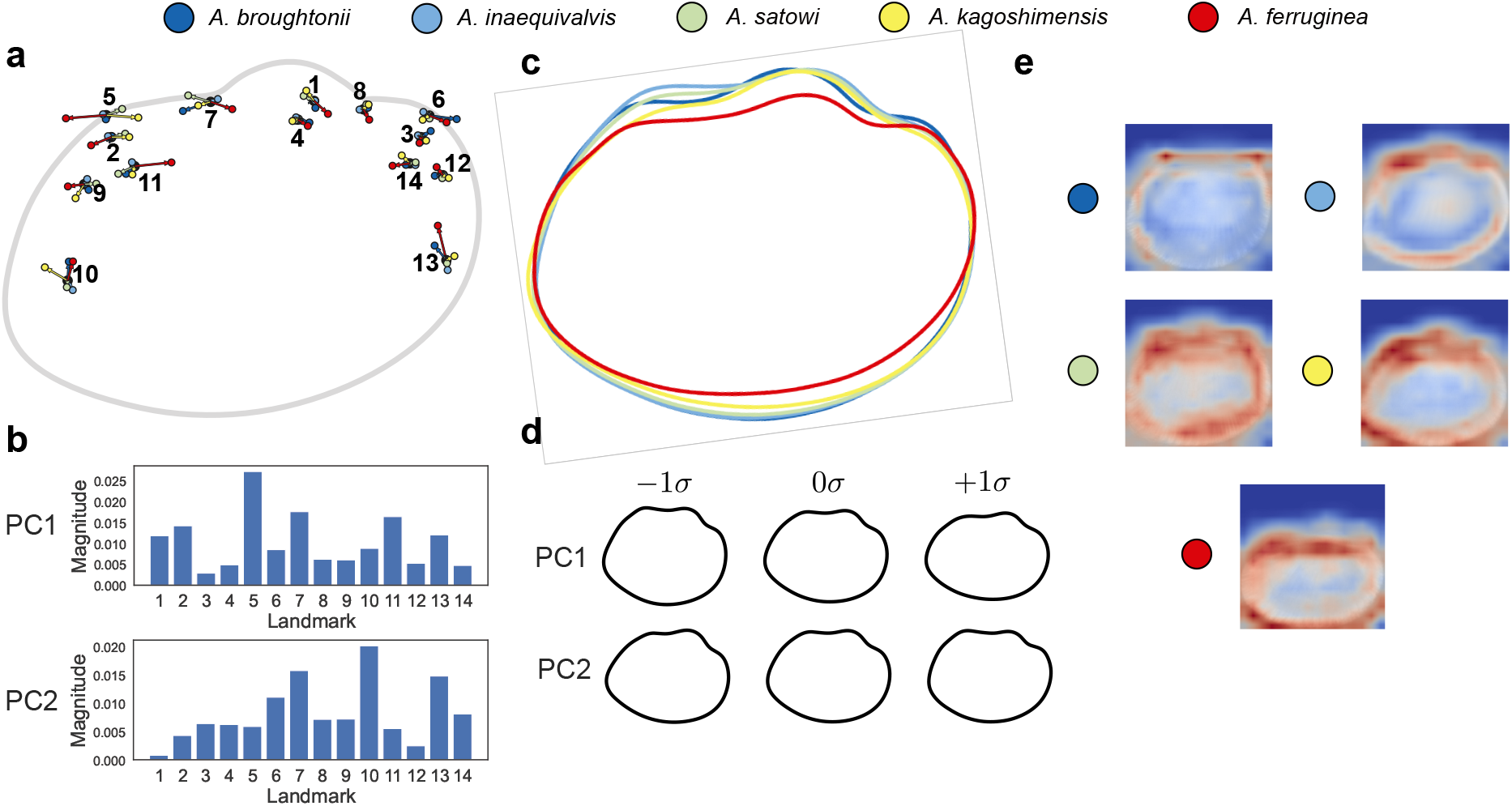
**a**, Displacement vectors showing the shape differences between species. Arrows originate from the overall mean shape and point toward each species’ mean landmark configuration, illustrating the direction and magnitude of the landmark displacement relative to the consensus shape. **b**, Principal component loadings for the landmark-based PCA, showing the contribution of each landmark (numbered 1-14) to PC1. **c**, Average shell contours reconstructed from the EFA coefficients for each species, showing interspecific shape differences based on the entire shell outline. **d**, Visualization of PC1 and PC2 mode variations in the EFA. PC1 and PC2 were derived by applying PCA to standardized elliptic Fourier descriptors (80 descriptors from coefficients up to the 20th harmonic). The shapes show morphological variations along the PC1 and PC2 axes. **e**, Saliency maps from Grad-CAM showing areas of importance for species identification by Morpho-VAE. The average heatmap for each species highlights the regions important for classification, with darker areas representing higher importance.

On the other hand, the mean contour for each species in the EFA is shown in Fig. 2c. *A. ferruginea* exhibits a shape with reduced height and increased width (Fig. 2c), resembling a shape with a high PC1 mode (Fig. 2d; PC1 +1*σ*), which is consistent with the fact that *A. ferruginea* has a high PC1 value (Fig. 1e). The mean contour of *A. inaequivalvis* showed a more protruding posterior dorsal corner (Fig. 2c; upper-left corner of the shell). This can be explained by *A. inaequivalvis* having a lower PC1 value than that of the other species.

The classification basis of Morpho-VAE was also visualized using Grad-CAM [39] (Fig. 2e), showing the regions judged to be important for identifying each species using deep learning. In *A. broughtonii*, although limited areas were emphasized, both sides of the dorsal corner were emphasized. In *A. inaequivalvis*, the posterior dorsal corner was emphasized, which is consistent with EFA results. The overall shell contour, specifically around the hinge line, was highlighted for all classes. Although Grad-CAM can visualize prominent regions (e.g., overall shell contour), this analysis fails to capture localized morphological features that can distinguish different classes. Other visualization frameworks that can quantitatively specify local regions that are important for classification and provide interpretable localized features are required.

### Reconstruction from the latent space of Morpho-VAE

While Fig. 2e visualizes the region highlighted by the classification module of Morpho-VAE, analyzing the VAE module is also possible. Morpho-VAE can reconstruct (i.e., generate) images from the latent space. This enables the visualization of how the VAE module of Morpho-VAE recognizes the morphological features in each class. Fig. 3a shows the results of the reconstructed images from the input data. The original image is first mapped into the latent space by the encoder, and then the reconstructed image is generated from the data point in the latent space by the decoder. In the reconstructed image, although the radial ribs are lost, the contour of the entire shell is well preserved, indicating that the latent space appropriately captures the global features of the shell morphology. The entire latent space can be visualized by generating images from the grid points in the latent space (Fig. 3c). The shell morphology continuously and smoothly changed in the latent space, suggesting that the model was not overfitted to a few samples. In addition, by setting a specific trajectory in the latent space, the morphological change from one class to another can be shown as morphing (Fig. 3b). For example, the morphology of *A. inaequivalvis* gradually transforms into that of *A. ferruginea*, emphasizing the morphological differences between them, such as the aspect ratio and appearance/disappearance of the umbo (Fig. 3d). However, these analyses are mainly based on the global features of morphology and do not extract local morphological features in an interpretable way.

**Fig. 3.**
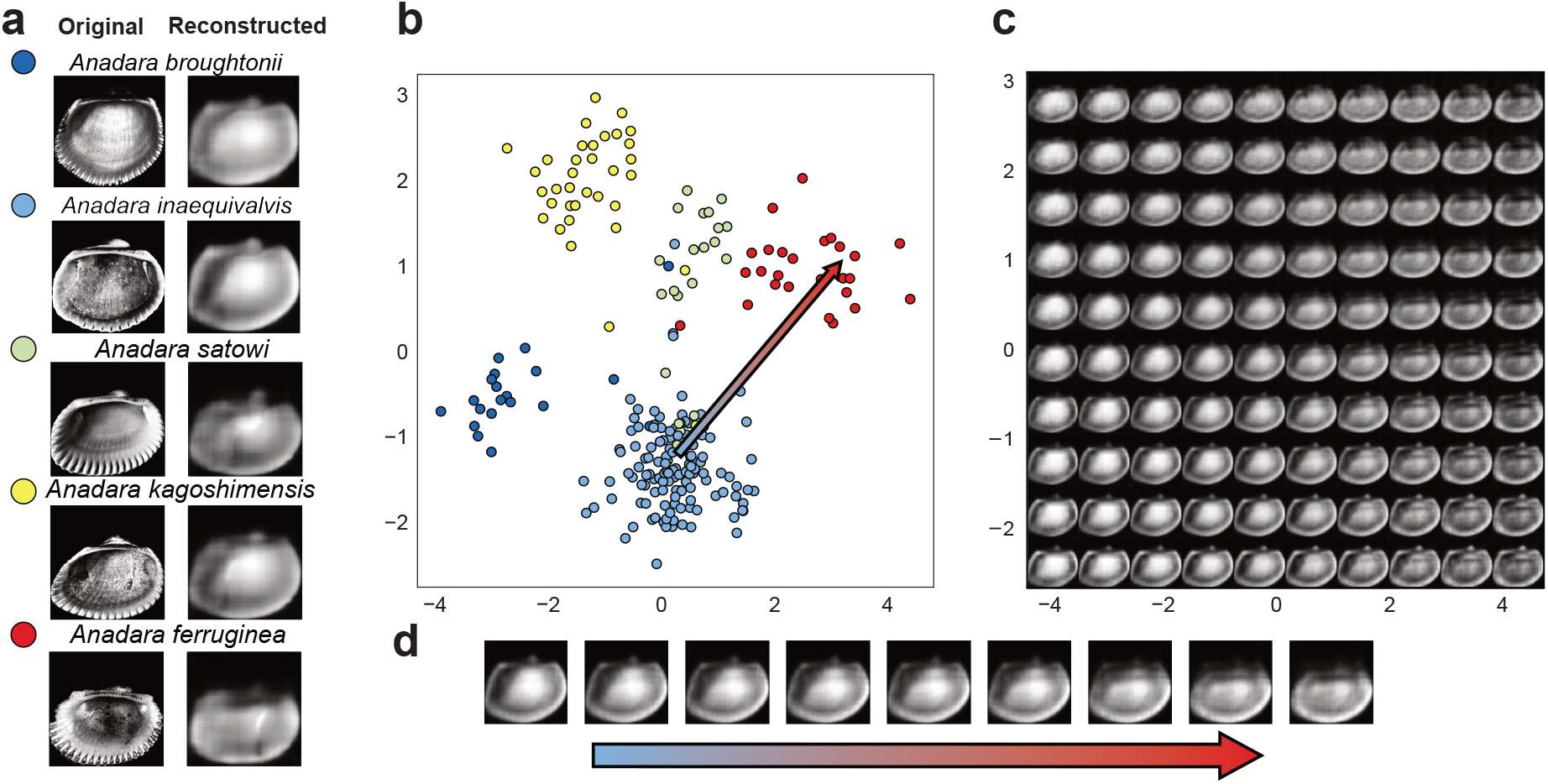
**a**, Illustration comparing the input data (left column) of the five *Anadara* species used in this study with the data reconstructed by Morpho-VAE (right column). **b**, Illustration of the Morpho-VAE latent space. A 2D latent space is visualized with the data points projected onto the space. The color of the points represents the species label. **c**, Images reconstructed by the decoder from the grid point coordinates in the latent space. The spatial range of the displayed plane corresponds to that shown in **b. d**, Morphing between *A. inaequivalvis* and *A. ferruginea* in latent space. The color of the point represents the species label, and the arrow represents the trajectory in the latent space.

### Patch partition to identify local morphological features

To analyze and quantify the influence of local morphological regions on species classification, we introduced a 4 *×* 4 grid partition for each image and systematically evaluated the contribution of each patch. This provides more interpretable local features that complement the global visualization offered by Grad-CAM.

To perform this analysis, we first constructed partially visible images in which only the selected one or more patches were visible (Figs. 4a-b top panels) and used them as input to a Morpho-VAE pre-trained on the original unmasked training dataset. From the confusion matrix that summarizes the results of class prediction, we computed the class-wise F1 score (see Methods) in a one-vs-rest manner. We then quantitatively evaluated the extent to which the F1 score increased by adding a specific patch. For instance, the F1 score of a partially visible image dataset in which only (X1, Y4), (X3, Y3), and (X4, Y2) patches were visible (Fig. 4a, top panel) was calculated as 0.10 for *A. inaequivalvis* using a pre-trained Morpho-VAE, and the score increased to 0.42 by adding (X2, Y3). For *A. ferruginea*, the score decreased from 0.45 to 0.44 (Fig. 4b). These results are summarized in the scatter plot in Fig. 4c, where a point above the diagonal line represents an improvement in the prediction performance.

**Fig. 4.**
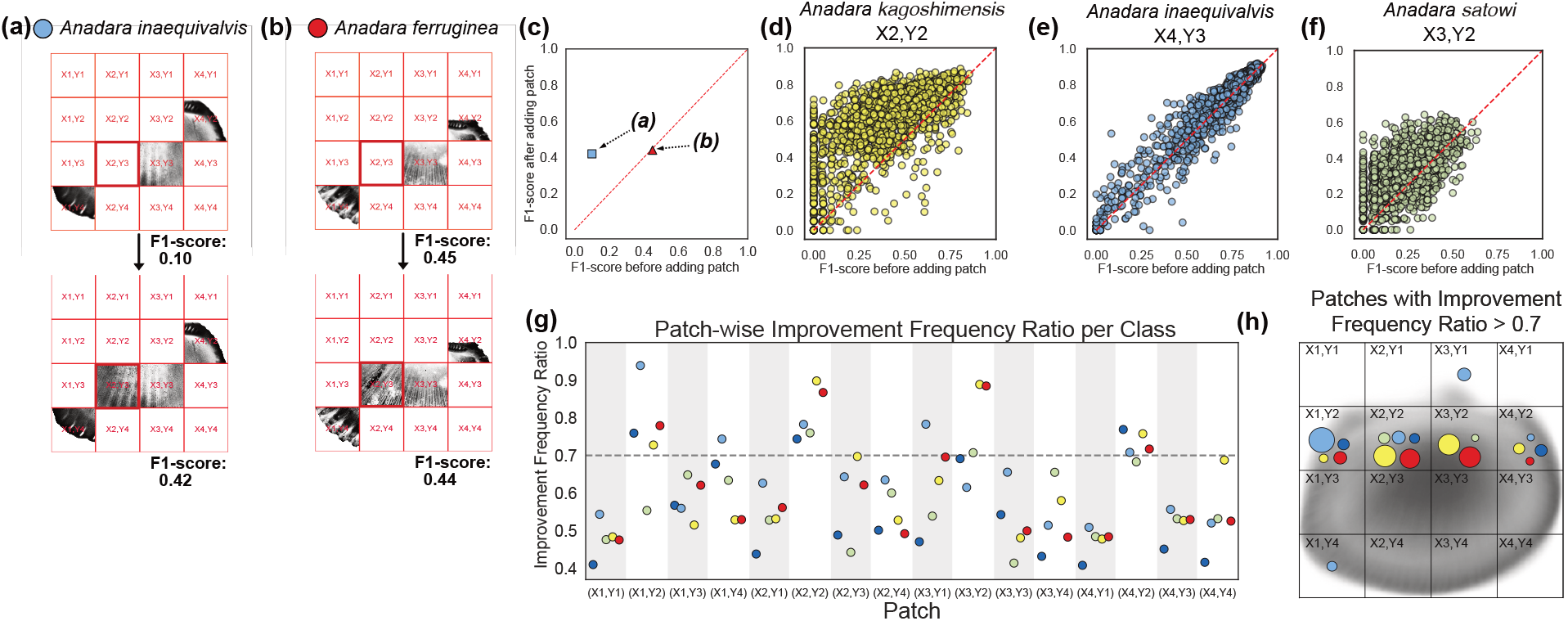
**a**, Effect of adding patch (X2, Y3) to patch combinations (X1, Y4), (X3, Y3), and (X4, Y2) in *Anadara inaequivalvis* (blue circle). An increase in the F1 score (from 0.10 to 0.42) was observed. **b**, Effect of adding patch (X2, Y3) to the patch combination (X1, Y4), (X3, Y3), and (X4, Y2) in *Anadara ferruginea* (red circle). A slight decrease in the F1 score (from 0.45 to 0.44) was observed. **c**, Scatter plot showing the F1 score before and after adding the patch (X2, Y3) for *Anadara inaequivalvis* (blue square) and *Anadara ferruginea* (red circle). The blue squares and red circles correspond to the examples shown in panels (a) and (b), respectively. **d-f**, Scatter plots showing the F1 score before and after adding target patches for **d**, *Anadara kagoshimensis* at (X2, Y2); **e**, *Anadara inaequivalvis* at (X4, Y3); and **f**, *Anadara satowi* at (X3, Y2). Each point represents a different combination of pre-existing patches before the target patch is added. **g**, Improvement Frequency Ratio (IFR) for each patch and species. The IFR is defined as the ratio of cases in which adding the target patch improves the F1 score. **h**, A distribution diagram showing patches and species where the IFR exceeds 0.7. The color of each circle corresponds to a species, and the size corresponds to the difference between each IFR and 0.7.

Fig. 4d shows the results of adding the patch (X2, Y2) for *A. kagoshimensis*. Each point represents one pre-existing patch combination evaluated with one pre-trained model before adding (X2, Y2). In this case, most points were above the diagonal line, indicating that adding the target patch (X2, Y2) was effective for predicting *A. kagoshimensis*. Fig. 4e illustrates the result for *A. inaequivalvis* with the target patch at (X4, Y3). This scatter plot suggests that the data points are distributed around the diagonal line, indicating that the addition of the (X4, Y3) patch does not increase the classification performance. Similarly, for *A. satowi* with the target patch at (X3, Y2) (Fig. 4f), the data points are located around the diagonal line, indicating that the target patch is not informative. In addition, all the F1 scores of the data points were below 0.65, indicating that the precise classification of *A. satowi* is difficult even when the entire image is visible.

Here, we define the Improvement Frequency Ratio (IFR) as the ratio of points above the diagonal line in the scatter plot in Figs. 4d-f, representing the fraction of cases in which the F1 score increased by adding the target patch to the number of all combinations of the pre-existing patch. Fig. 4g summarizes the IFR for each patch and species computed by performing a before-and-after comparison of the F1 score for all combinations. This analysis revealed that the contribution of adding individual patches was limited, with most IFR values falling below a heuristically determined threshold, that is, the IFR was 0.7 (Fig. 4g). Despite this general trend, the analysis successfully identified several patches that were highly effective in classifying specific species. The most notable finding was that the hinge line corresponding to the patches (X1–X4, Y2) was identified as a key determinant for all species. Specifically, the local feature around (X1, Y2) is important for *A. inaequivalvis*, whereas characteristic features are localized at (X2, Y2) and (X3, Y2) for *A. kagoshimensis* and *A. ferruginea*. Besides the hinge line, the umbo (X3, Y1) and anterior ventral margin (i.e., bottom left corner of the shell) (X1, Y4) are important for *A. inaequivalvis*. These morphologically informative localized regions are illustrated in Fig. 4h.

## DISCUSSION

In this study, we applied the deep-learning-based morphological analysis using Morpho-VAE to the shell shape classification of *Anadara* species. The proposed method does not require defining landmarks or contour lines, offering a practical and versatile framework for morphological analysis. A comparative analysis with well-established methods showed that Morpho-VAE significantly outperformed the landmark-based approaches and EFA in a species classification task, demonstrating the advantages of deep-learning-based methods in capturing richer and more discriminative morphological features. This reflects the model’s ability to process the entire image and its supervised latent space construction, in contrast to the unsupervised latent spaces of EFA or landmark-based methods.

Our patch analysis identified several important morphological traits for classification. The most notable region was the hinge line (Fig. 4h; (X1–X4, Y2)). While the importance of the hinge line is consistent with the positions of the landmarks (Fig. 1b and Fig. 2a), in Morpho-VAE, the top left corner of the shell (X1, Y2) was identified as the most important determinant of *A. inaequivalvis*, which was not the case in the landmark analysis (see the length of the light blue arrow at landmarks 2, 5, 7, and 11). Morpho-VAE also found key morphological regions at the bottom left corner (X1, Y4) and the umbo position (X3, Y1) to classify *A. inaequivalvis*; however, no landmarks were defined in these regions. These findings imply that Morpho-VAE is useful for identifying the morphological features that can be overlooked by conventional methods. Our method may also help guide the placement of landmarks and can thus be used as a complement to the landmark method.

Recently, numerous studies on deep-learning-based morphological analysis have been conducted [31, 33–35, 37]. While approaches such as metric learning [33, 34] focus on reflecting phylogenetic relationships in the latent space, Morpho-VAE emphasizes classification performance and interpretability through its generative decoder. These approaches are complementary, depending on the research objectives. While this study used Morpho-VAE, when seeking to more directly reflect bivalve phylogenetic information or incorporate phylogenetic tree data, analyses using metric learning based on phylogenetic pairs/triplets (or combining VAE with distance learning) to reorganize the latent space according to phylogenetic relationships may be effective.

A clear limitation of this study is that the reconstructed images generated by Morpho-VAE (Fig. 3a) fail to capture fine texture information, such as radial ribs on the shell surface, which are traditional diagnostic traits [40, 41]. Consequently, the overall image appears blurred. This is attributed to the fact that the high-frequency components of the image, that is, fine structures, are easily lost during dimensionality reduction, because the positions or even the presence/absence of these ribs can largely fluctuate among images. For instance, the position or number of radial ribs can change among samples, even within the same species [41].

In summary, Morpho-VAE provided an improved low-dimensional representation of inter-specific variation in *Anadara* shells, compared with landmark-based approaches and EFA. Combined with the patch-based analysis, our approach identified the primary determinants of image-based discrimination as morphological features, including the hinge line, its associated internal structures, and regions such as the umbo and ventral margin. The present pipeline provides a powerful tool for discovering interpretable morphological traits to identify interspecific differences. The extension to incorporate multimodal input such as images with different resolutions and scales to better capture fine sculptures (e.g., radial ribs) needs to be addressed in the future.

## METHODS

### Image preprocessing

We collected shells of five *Anadara* species: *Anadara kagoshimensis* (Tokunaga, 1906), *Anadara inaequivalvis* (Bruguière, 1789), *Anadara satowi* (Dunker, 1882), *Anadara broughtonii* (Schrenck, 1867), and *Anadara ferruginea* (Reeve, 1844). In total, 230 left valves were used in this study, comprising 20 *A. broughtonii*, 127 *A. inaequivalvis*, 22 *A. satowi*, 35 *A. kagoshimensis*, and 26 *A. ferruginea*. All samples were preserved at The University Museum, The University of Tokyo. The details and distribution of the samples are presented in Supplementary Table 1. All samples were photographed using a digital camera to capture the shell outlines. To ensure consistency for the *Anadara* genus, which possesses hinge teeth, the teeth were aligned parallel to the camera’s horizontal axis, and the shells were secured with clay to maintain their surfaces parallel to the focal plane (SFig. 5).

After photographing the shells, we performed a series of preprocessing steps to ensure image uniformity and improve the analysis accuracy. First, we extracted the outlines of the shells from the original images using Adobe Photoshop (SFig. 6b). We then trimmed the image and resized it to 256 *×* 256 pixels, setting the lowest vertical point of each shell as the reference point (SFig. 6c). The images were then converted to grayscale (SFig. 6d). However, in conventional photographs, the pixel values were concentrated in the higher range (with 0 corresponding to the background), as shown in the histogram in SFig. 6d. Because subsequent analyses required the full spectrum of pixel values, we applied contrast-limited adaptive histogram equalization (CLAHE) to redistribute the pixel values across the entire range, enhancing the contrast and ensuring better feature extraction (SFig. 6e).

### Geometric morphometrics based on landmarks

For the landmark-based analysis, the internal features of the same 230 left valves were photographed using the setup described above. On each image, 14 landmarks were digitized on the internal shell features using ImageJ [42] (Fig. 1). The *x−y* coordinates for each sample were compiled and imported into MorphoJ [43]. Generalized Procrustes analysis (GPA) was conducted to rotate, scale, and translate the landmark positions, yielding Procrustes coordinates. A covariance matrix was then generated, followed by PCA.

To determine significant morphological traits in *Anadara*, interspecific differences were quantitatively evaluated and visualized using PCA scores. We computed the mean shape for each species, plus the collective mean shape for every specimen. These mean PCA scores were inversely transformed back into the Procrustes coordinate space to reconstruct the landmark positions. Finally, displacement vectors were calculated as the difference between the various average shapes, and the average shape was visualized as arrows indicating the direction and magnitude of morphological variation (Fig. 2a).

### EFA on the shell shape

For EFA, we used the same dataset of 230 left valves (20 *A. broughtonii*, 127 *A. inaequivalvis*, 22 *A. satowi*, 35 *A. kagoshimensis*, and 26 *A. ferruginea*). The mask images generated during preprocessing were used to extract contour *x − y* coordinates via OpenCV. We used the Python library `pyefd` to derive elliptic Fourier descriptors. The descriptors were standardized to account for the size (scaling the semi-major axis of the first harmonic) and orientation (aligning the phases of the semi-major axis).

In this study, the contours were approximated using coefficients up to the 20th harmonic, resulting in 80 descriptors (4 coefficients *×* 20 harmonics) per shell. PCA was applied to these standardized coefficients to identify the primary patterns of interspecific variation. The mean Fourier coefficients for each species were computed and subjected to inverse Fourier transformation to reconstruct the representative average contour shapes. In addition, individual contours were reconstructed and overlaid to evaluate intraspecific variation. This quantitative approach provided a robust characterization of interspecific shape differences based on the entire shell outline.

### Morpho-VAE parameter tuning and architecture details

In this study, we used Morpho-VAE [37] to extract low-dimensional morphological features from *Anadara* shell images for species classification. The key advantage of Morpho-VAE lies in its ability to automatically extract morphological features directly from raw image data, eliminating the need for manual landmark annotations. This reduces manual effort and the potential for human error and enables the detection of subtle morphological details that are often overlooked by conventional methods of analysis.

In previous studies [37], the input images were binary, which limited the ability to capture morphological features such as coloration and sculpture patterns. To address this issue, we replaced the binary images with grayscale images, where all pixel values are represented as real numbers, enabling a more detailed morphological analysis.

Morpho-VAE comprises three main components (SFig. 8):

1. The encoder transforms 256 *×* 256 images into a low-dimensional latent space.
2. The decoder reconstructs the input image from the latent representation of the image.
3. The classification module branches off from the latent space to predict the species label.

The encoder comprises convolutional layers combined with MaxPooling layers, with the final layer outputting the mean and log variance of an *N*_*z*_-dimensional latent space. The decoder comprises convolutional and upsampling layers that generate images with pixel values in the range [0,1]. In parallel, the classification module applies the Softmax function to output species predictions. Unless otherwise noted, Morpho-VAE analyses in this study were performed with a three-dimensional latent space (*N*_*z*_ = 3), whereas Fig. 1c shows a two-dimensional latent space for visualization.

### Parameter tuning of Morpho-VAE

The hyperparameters of Morpho-VAE, including the number of layers, number of filters per layer, activation function, and optimization function, were tuned using Optuna [44]. Following previous studies [37], we optimized the number of layers within the range of 1–9 and tuned the number of filters per layer within the range of 16–128. The activation function was selected from ReLU, sigmoid, and tanh, whereas the optimization function was selected from stochastic gradient descent (SGD), Adam, and RMSprop. During optimization, we evaluated 500 hyperparameter sets, training each model for 1000 epochs to determine the optimal configuration that minimized the loss functions. To evaluate model performance while mitigating overfitting, we adopted a two-stage, group-aware data partitioning strategy. In the first stage, we split the data into training and test subsets using GroupShuffleSplit (train size = 0.67, test size = 0.33) so that images sharing the same specimen tag were not assigned to both subsets. To preserve class balance, this group-wise split was performed separately within each class and then concatenated across classes. In the second stage, the training subset was further divided into training and validation subsets using the same procedure (train size = 0.75, validation size = 0.25). This produced three non-overlapping datasets (Training, Validation, and Test), prevented specimen-level data leakage, and enabled unbiased model evaluation.

Each trial was evaluated based on a weighted loss function comprising the following components:

1. Reconstruction Loss (*E*_*Rec*_) This loss function is defined as the mean squared pixel-wise error between the input vector (**x**^(*i*)^ *∈* ℝ^*D*^) and reconstructed vector (**x**′^(*i*)^ *∈* ℝ^*D*^), averaged over all pixels (*D* = 256^2^ = 65, 536) and all samples (*N*)

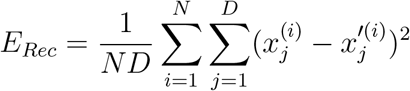
2. Classification Loss (*E*_*C*_) This loss function is defined as the categorical cross-entropy between the predicted class probability vector **y**′^(*i*)^, obtained from the classification module with latent variable **z**^(*i*)^ as input, and the true one-hot label vector **y**^(*i*)^. For each sample, the cross-entropy is summed over all classes, and then averaged over all samples *N* :

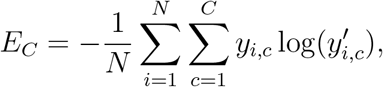

where *C* = 5 in this study.
3. Regularization Loss (*E*_*Reg*_) This loss term constrains the latent representation so that the distribution of the latent variable **z** does not deviate excessively from a unit Gaussian. It penalizes both large absolute values of the mean vector ***μ*** and large variances ***σ***^2^ as follows:

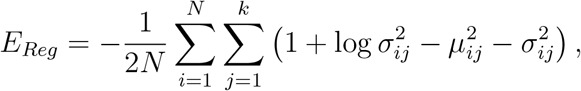

where *k* is the dimension of the latent variable **z**.

The total loss function is defined as follows:

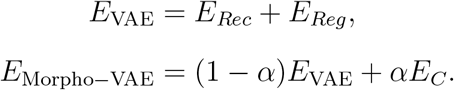

We used *α* = 0.9 for the analyses reported in the main text; this value was selected from a Pareto-based trade-off between reconstruction and classification losses on validation data over a grid of *α* values (SFig. 7), as summarized later in this section.

### Optimized model configuration

After optimization, the encoder was configured with eight layers, with filter sizes set to 64, 128, 32, 80, 96, 64, 128, and 128 from the input layer. The activation function used was tanh, and the optimization function was RMSprop. The decoder also comprised eight layers, with filter sizes set to 128, 128, 64, 96, 80, 32, 128 and 64, from the layer closest to the latent variable. Because the input images were grayscale (values in the range [0,1]), a sigmoid activation function was used in the final decoder layer instead of tanh to ensure that the reconstructed images remained within the same range.

We also optimized the value of *α*, which is a critical hyperparameter that significantly influences the performance of the Morpho-VAE model. The value of *α* controls the weight balance between the reconstruction and classification losses, playing a crucial role in adjusting the trade-off between reconstruction accuracy and classification performance during the learning process of the model. For the optimization experiments, we set 11 levels of *α* values from 0 to 1.0 in 0.1 increments (0, 0.1, 0.2, 0.3, 0.4, 0.5, 0.6, 0.7, 0.8, 0.9, 1.0), and executed experiments for each value with 10 random seeds. In the experimental configuration, the model architecture comprised eight convolutional layers with filter numbers of 64, 128, 32, 80, 96, 64, 128, and 128 for each layer. We used the RMSprop optimizer and tanh activation function for optimization, and training was executed for up to 1000 epochs with a batch size of 10. During training, we monitored the validation loss and stopped training early when further improvement was no longer observed, so some runs terminated before reaching the maximum number of epochs.

The performance evaluation for each *α* value was conducted using *E*_*Rec*_ and *E*_*C*_.

In the optimization process, we statistically analyzed the experimental results for each *α* value across 10 random seeds. First, we computed the *E*_*C*_ and *E*_*Rec*_ for each *α* value over 10 validation runs (one per random seed), and calculated *E*_Morpho*−*VAE_ for each *α* (SFig. 7a). We did not include *E*_*Reg*_ in the optimization process, because the value of *E*_*Reg*_ was too small to affect the result. Then, to quantitatively evaluate the trade-off between the reconstruction and classification errors on the validation data, we aggregated *E*_*C*_ and *E*_*Rec*_ on the validation data for each trial using *α*, and calculated their means and standard deviations. These results were plotted on a two-dimensional plane, and Pareto efficiency was used to identify the Pareto front (SFig. 7b). The standard deviations are shown as error bars, and the Pareto-optimal points are highlighted with annotations of the corresponding *α* values. From this optimization process, we selected an *α* value of 0.9 as the optimal value, demonstrating a good balance between reconstruction accuracy and classification performance.

### Patch-based analysis

To evaluate which local morphological features are important for species classification, we applied a patch-based analysis to the Morpho-VAE model. First, we selected one or more patches from a set of 16 patches, displayed only the image corresponding to the selected patch (i.e., unselected patches were masked), and used this image as input for prediction. Predictions across all species produced a confusion matrix, which was used to evaluate the species classification performance. Owing to the sample size imbalance among species, we evaluated the patch effects using class-wise F1 scores computed in a one-vs-rest manner. For each confusion matrix associated with a given condition (with vs. without a target patch) and test seed, we derived true positive (TP), false positive (FP), false negative (FN), and true negative (TN) for each class *c*, and then calculated precision 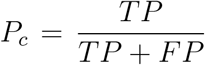, recall 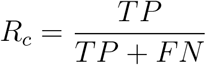, and F1 score 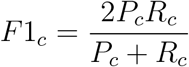. The F1 score is less sensitive to class priors than accuracy and thus provides a more reliable comparison under class imbalance.

We then quantified the change in the F1 score (ΔF1) before and after adding a single patch as the unit effect. For a given patch *P* and test seed, we calculated the F1 score from two matched conditions: (i) mask images without a patch *P*, and (ii) mask images with a patch *P*. For each condition, we computed F1 score *F*1_before_ (case (i)) and *F*1_after_ (case (ii)).

Figs. 4a-b show a before-after comparison of the F1 score for *A. ferruginea* and *A. inaequivalvis* with the same target patch at (*X*2,*Y* 3). In every panel, the upper row shows the image without the target patch, and the lower row shows the image with the target patch. Fig. 4c shows a scatter plot of the F1 score before and after adding the target patch. Each dot represents one mask-combination/model condition. This visualization suggests that the farther a point is above the diagonal line, the greater the improvement in prediction performance achieved by adding the target patch. Figs. 4d-f are scatter plots illustrating the results of all before-after comparisons for *A. kagoshimensis, A. inaequivalvis*, and *A. satowi* at (X2, Y2), (X4, Y3), and (X3, Y2), respectively.

We then computed the change in the F1 score (ΔF1) as ΔF1 = *F*1_after_ - *F*1_before_. Aggregating these changes across all models and mask pairs that contain patch *P*, we obtained the IFR, which is the fraction of positive ΔF1 values in all combinations that include patch *P*. If the IFR was greater than 70%, we considered that patch *P* was effective for species classification. Fig. 4g shows a summary of the IFR for all patches in all models. The gray dotted line indicates a threshold of 70%. Fig. 4h illustrates all patches where the IFR is greater than 70% and the point size corresponds to the difference between the IFR value and the threshold.

## ACKNOWLEDGMENTS

We thank Hideaki Sato for the helpful discussions. This research was supported in part by JSPS KAKENHI (25H01364 and 25K07242 to N.S.; 22K21344 and 23K27164 to C.F.). Masato Tsutsumi was supported by Takeda Science Foundation.

