## Supplementary Figures and Tables for "Decoding *Anadara* shell morphology with deep learning"

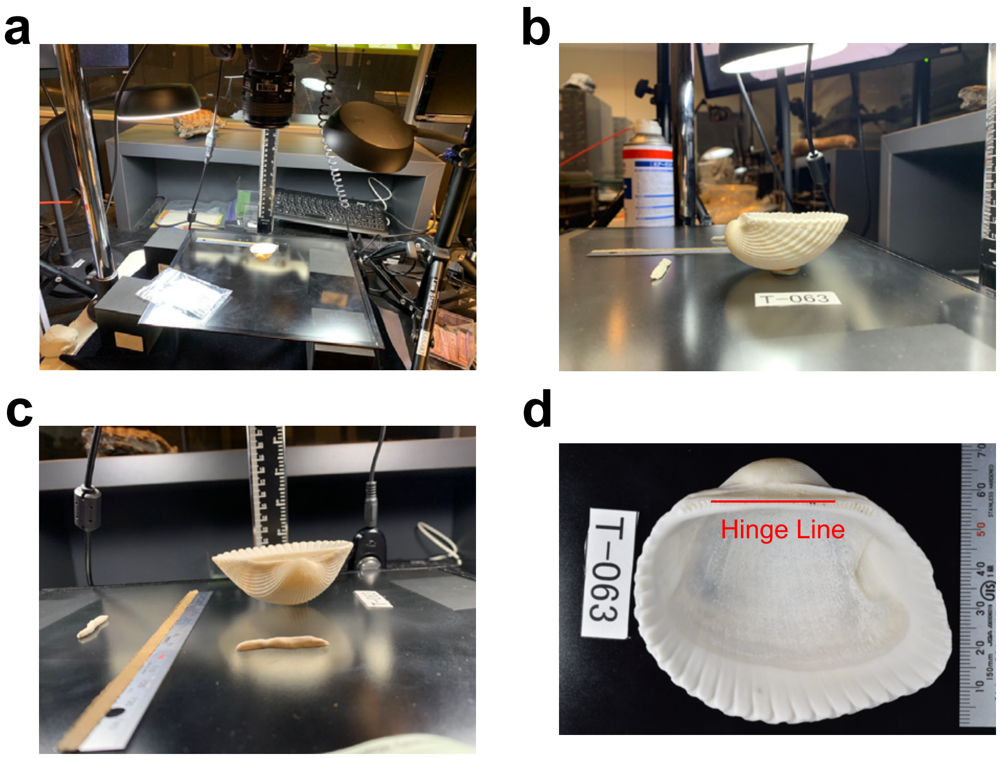


SFig. 1. **a**, Photograph of the photographing platform, digital camera, and lighting. This study was conducted at the University Museum, the University of Tokyo. **b**, Photograph of a shell fine-tuned with paper clay so that its surface was level with the photographing platform. **c**, Photograph of a shell being adjusted using paper clay to level it in the left and right directions. **d**, Photograph of *Anadara inaequivalvis* taken using the aforementioned protocol. A ruler was placed to show the scale. A piece of paper with an ID was placed in the upper left corner to distinguish the photographs. The red line represents the hinge line, which serves as a horizontal reference.

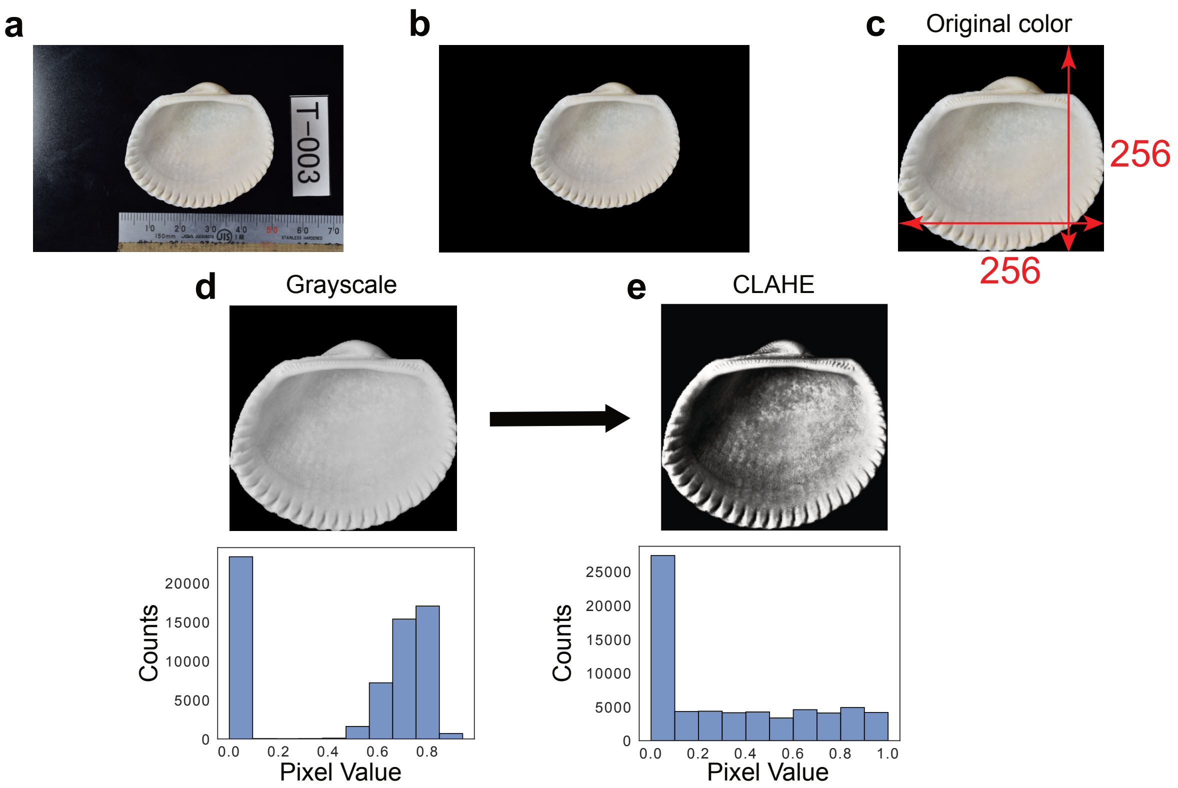


SFig. 2. Processing workflow applied to *Anadara* shell photographs for analysis. **a** Original photograph acquired on the imaging platform, showing the specimen, scale bar, and sample ID. **b** Extracted outline (mask) produced using Adobe Photoshop to isolate the shell contour from the background. **c** Aligned and cropped image: images were trimmed and vertically aligned using the lowest shell point as a reference and then resized to 256 $\times$ 256 pixels. **d** Grayscale conversion and corresponding pixel-value histogram, illustrating the original brightness concentration toward higher values. **e** Contrast-enhanced image after applying contrast-limited adaptive histogram equalization (CLAHE), producing a more uniform pixel distribution used as the final input for downstream analyses.

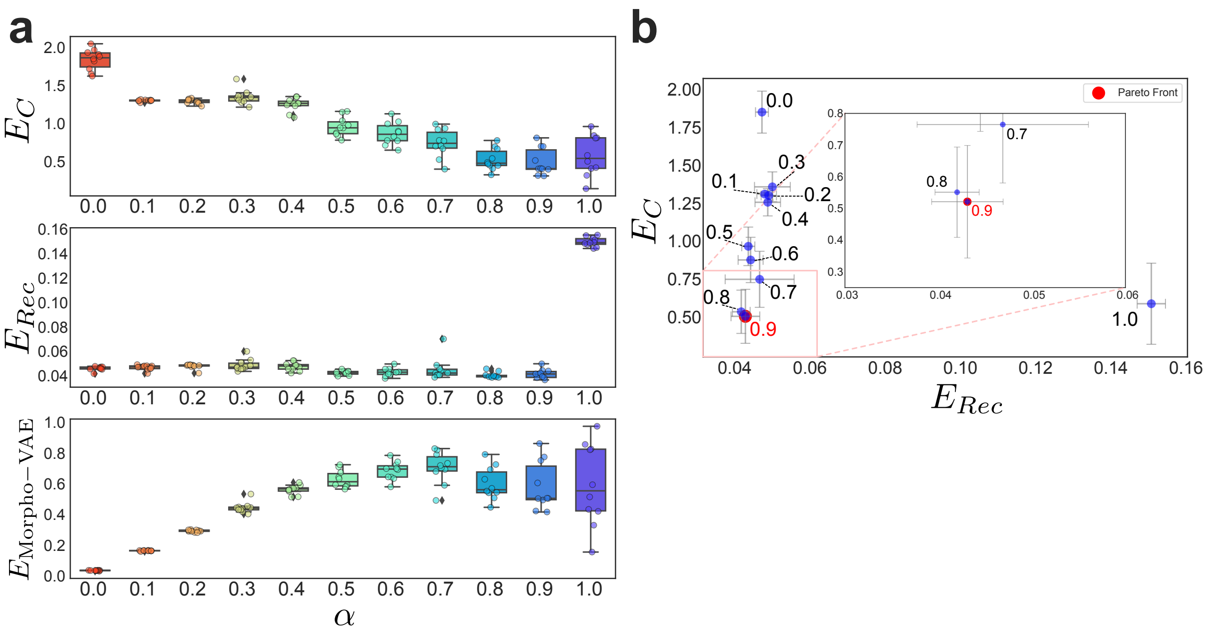


SFig. 3. **a**, Reconstruction loss ($E_{Rec}$), classification loss ($E_C$), and combined Morpho-VAE loss ($E_{\text{Morpho-VAE}}$) computed across validation data points for each $\alpha$ value (0.0-1.0). Error bars indicate the standard deviation across random seeds/trials. **b**, Pareto front analysis plotting $E_{Rec}$ against $E_C$ for each $\alpha$. Error bars represent the standard deviations across trials. Each point on the Pareto front is highlighted with the corresponding $\alpha$ annotation, indicating that $\alpha = 0.9$ represents the optimal value in the trade-off between the reconstruction and classification performances.

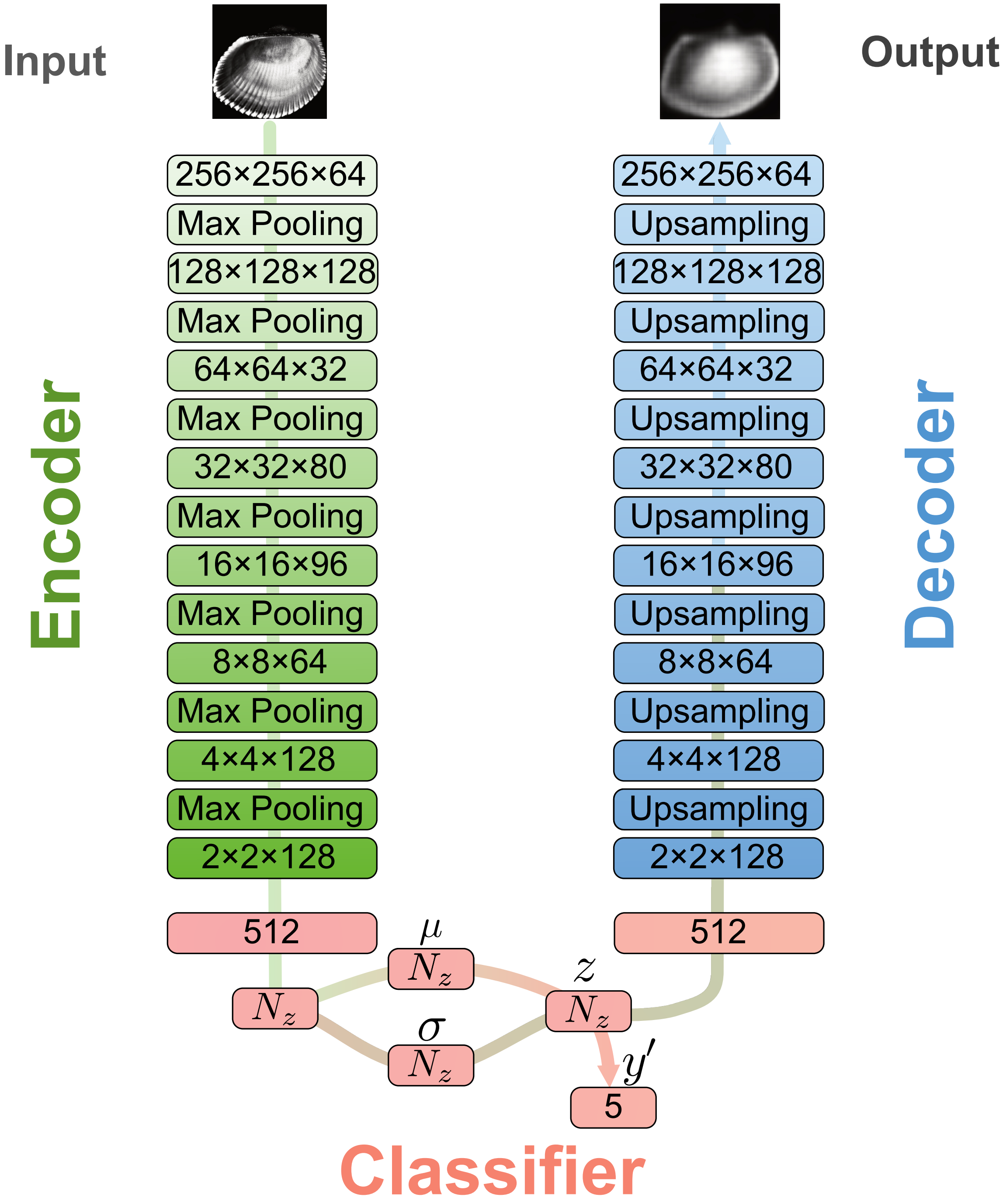


SFig. 4. Schematic of the Morpho-VAE model used for the *Anadara* shell images. The input images were processed using a convolutional encoder to form a compressed latent space $z$ (characterized by $\mu$ and $\sigma$). This latent representation is used for two parallel tasks: (1) image reconstruction via the decoder and (2) classification into five categories ($y'$) via the classifier branch. The numbers in the blocks indicate the tensor dimensions of each layer of the deep neural network.

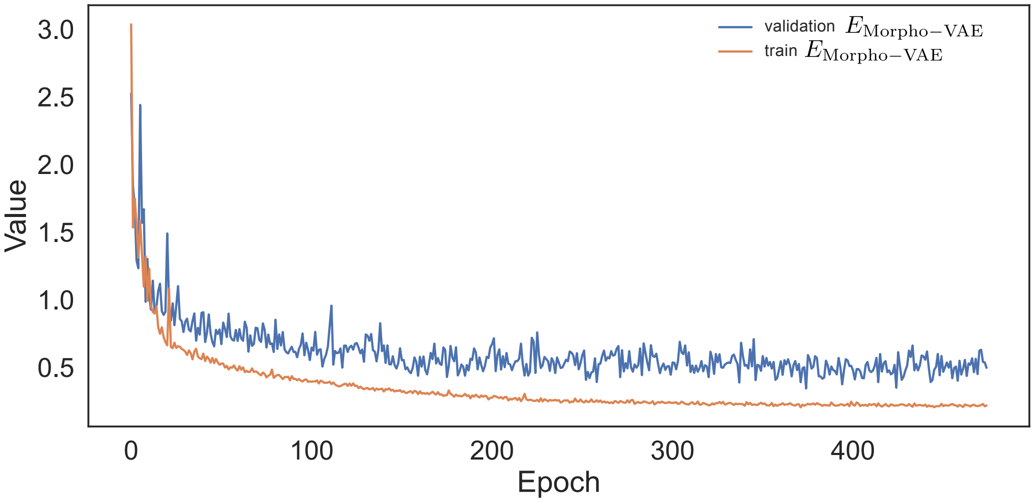


SFig. 5. Learning curves of the Morpho-VAE model for the *Anadara* shell images. The plot shows the transition of the training (orange) and validation (blue) loss ($E_{\text{Morpho-VAE}}$). Although the maximum number of training epochs was set to 1000, training terminated at around 400 epochs because of EarlyStopping; therefore, only the observed epoch range is shown.

STable. I: Complete list of specimens examined in this study, grouped by species.

| No. | Scientific Name | Locality |
|---|---|---|
| 1 | *Anadara broughtonii* | Fish market at Tokyo, Japan |
| 2 | *Anadara broughtonii* | Fish market at Tokyo, Japan |
| 3 | *Anadara broughtonii* | Fish market at Tokyo, Japan |
| 4 | *Anadara broughtonii* | Fish market at Tokyo, Japan |
| 5 | *Anadara broughtonii* | Yamaguchi Bay, Yamaguchi, Japan |
| 6 | *Anadara broughtonii* | Yamaguchi Bay, Yamaguchi, Japan |
| 7 | *Anadara broughtonii* | Yamaguchi Bay, Yamaguchi, Japan |
| 8 | *Anadara broughtonii* | Yamaguchi Bay, Yamaguchi, Japan |
| 9 | *Anadara broughtonii* | Yamaguchi Bay, Yamaguchi, Japan |
| 10 | *Anadara broughtonii* | Yamaguchi Bay, Yamaguchi, Japan |
| 11 | *Anadara broughtonii* | Yamaguchi Bay, Yamaguchi, Japan |
| 12 | *Anadara broughtonii* | Yamaguchi Bay, Yamaguchi, Japan |
| 13 | *Anadara broughtonii* | Yamaguchi Bay, Yamaguchi, Japan |
| 14 | *Anadara broughtonii* | Yamaguchi Bay, Yamaguchi, Japan |
| 15 | *Anadara broughtonii* | Yamaguchi Bay, Yamaguchi, Japan |
| 16 | *Anadara broughtonii* | Yamaguchi Bay, Yamaguchi, Japan |
| 17 | *Anadara broughtonii* | Yamaguchi Bay, Yamaguchi, Japan |
| 18 | *Anadara broughtonii* | Yamaguchi Bay, Yamaguchi, Japan |
| 19 | *Anadara broughtonii* | Yamaguchi Bay, Yamaguchi, Japan |
| 20 | *Anadara broughtonii* | Yamaguchi Bay, Yamaguchi, Japan |
| 21 | *Anadara ferruginea* | Off Kushimoto, Wakayama, Japan |
| 22 | *Anadara ferruginea* | Off Kushimoto, Wakayama, Japan |
| 23 | *Anadara ferruginea* | Off Kushimoto, Wakayama, Japan |
| 24 | *Anadara ferruginea* | Off Kushimoto, Wakayama, Japan |
| 25 | *Anadara ferruginea* | Off Kushimoto, Wakayama, Japan |
| 26 | *Anadara ferruginea* | Off Kushimoto, Wakayama, Japan |
| 27 | *Anadara ferruginea* | Off Kushimoto, Wakayama, Japan |

STable. I: Complete list of specimens examined in this study, grouped by species.

| No. | Scientific Name | Locality |
|---|---|---|
| 28 | *Anadara ferruginea* | Off Kushimoto, Wakayama, Japan |
| 29 | *Anadara ferruginea* | Off Kushimoto, Wakayama, Japan |
| 30 | *Anadara ferruginea* | Off Kushimoto, Wakayama, Japan |
| 31 | *Anadara ferruginea* | Off Kushimoto, Wakayama, Japan |
| 32 | *Anadara ferruginea* | Off Kushimoto, Wakayama, Japan |
| 33 | *Anadara ferruginea* | Off Kushimoto, Wakayama, Japan |
| 34 | *Anadara ferruginea* | Off Kushimoto, Wakayama, Japan |
| 35 | *Anadara ferruginea* | Off Kushimoto, Wakayama, Japan |
| 36 | *Anadara ferruginea* | Off Mishima Island, Yamaguchi, Japan |
| 37 | *Anadara ferruginea* | Off Mishima Island, Yamaguchi, Japan |
| 38 | *Anadara ferruginea* | Off Suhara, Wakayama, Japan |
| 39 | *Anadara ferruginea* | Off Suhara, Wakayama, Japan |
| 40 | *Anadara ferruginea* | Off Suhara, Wakayama, Japan |
| 41 | *Anadara ferruginea* | Off Suhara, Wakayama, Japan |
| 42 | *Anadara ferruginea* | Off Suhara, Wakayama, Japan |
| 43 | *Anadara ferruginea* | Off Suhara, Wakayama, Japan |
| 44 | *Anadara ferruginea* | Off Jogashima Island, Kanagawa, Japan |
| 45 | *Anadara ferruginea* | Off Jogashima Island, Kanagawa, Japan |
| 46 | *Anadara ferruginea* | Off Jogashima Island, Kanagawa, Japan |
| 47 | *Anadara inaequivalvis* | Katase-Higashi Beach, Kanagawa, Japan |
| 48 | *Anadara inaequivalvis* | Katase-Higashi Beach, Kanagawa, Japan |
| 49 | *Anadara inaequivalvis* | Katase-Higashi Beach, Kanagawa, Japan |
| 50 | *Anadara inaequivalvis* | Katase-Higashi Beach, Kanagawa, Japan |
| 51 | *Anadara inaequivalvis* | Zushi Beach, Kanagawa, Japan |
| 52 | *Anadara inaequivalvis* | Zushi Beach, Kanagawa, Japan |
| 53 | *Anadara inaequivalvis* | Zushi Beach, Kanagawa, Japan |
| 54 | *Anadara inaequivalvis* | Zushi Beach, Kanagawa, Japan |

STable. I: Complete list of specimens examined in this study, grouped by species.

| No. | Scientific Name | Locality |
|---|---|---|
| 55 | *Anadara inaequivalvis* | Zushi Beach, Kanagawa, Japan |
| 56 | *Anadara inaequivalvis* | Funakoshi, Akita, Japan |
| 57 | *Anadara inaequivalvis* | Nijino-matsubara, Saga, Japan |
| 58 | *Anadara inaequivalvis* | Nijino-matsubara, Saga, Japan |
| 59 | *Anadara inaequivalvis* | Tsuyazaki, Fukuoka, Japan |
| 60 | *Anadara inaequivalvis* | Tsuyazaki, Fukuoka, Japan |
| 61 | *Anadara inaequivalvis* | Tsuyazaki, Fukuoka, Japan |
| 62 | *Anadara inaequivalvis* | Fukiagehama Beach, Kagoshima, Japan |
| 63 | *Anadara inaequivalvis* | Fukiagehama Beach, Kagoshima, Japan |
| 64 | *Anadara inaequivalvis* | Fukiagehama Beach, Kagoshima, Japan |
| 65 | *Anadara inaequivalvis* | Fukiagehama Beach, Kagoshima, Japan |
| 66 | *Anadara inaequivalvis* | Fukiagehama Beach, Kagoshima, Japan |
| 67 | *Anadara inaequivalvis* | Fukiagehama Beach, Kagoshima, Japan |
| 68 | *Anadara inaequivalvis* | Fukiagehama Beach, Kagoshima, Japan |
| 69 | *Anadara inaequivalvis* | Fukiagehama Beach, Kagoshima, Japan |
| 70 | *Anadara inaequivalvis* | Fukiagehama Beach, Kagoshima, Japan |
| 71 | *Anadara inaequivalvis* | Fukiagehama Beach, Kagoshima, Japan |
| 72 | *Anadara inaequivalvis* | Fukiagehama Beach, Kagoshima, Japan |
| 73 | *Anadara inaequivalvis* | Fukiagehama Beach, Kagoshima, Japan |
| 74 | *Anadara inaequivalvis* | Fukiagehama Beach, Kagoshima, Japan |
| 75 | *Anadara inaequivalvis* | Fukiagehama Beach, Kagoshima, Japan |
| 76 | *Anadara inaequivalvis* | Fukiagehama Beach, Kagoshima, Japan |
| 77 | *Anadara inaequivalvis* | Fukiagehama Beach, Kagoshima, Japan |
| 78 | *Anadara inaequivalvis* | Fukiagehama Beach, Kagoshima, Japan |
| 79 | *Anadara inaequivalvis* | Fukiagehama Beach, Kagoshima, Japan |
| 80 | *Anadara inaequivalvis* | Fukiagehama Beach, Kagoshima, Japan |
| 81 | *Anadara inaequivalvis* | Fukiagehama Beach, Kagoshima, Japan |

STable. I: Complete list of specimens examined in this study, grouped by species.

| No. | Scientific Name | Locality |
|---|---|---|
| 82 | *Anadara inaequivalvis* | Fukiagehama Beach, Kagoshima, Japan |
| 83 | *Anadara inaequivalvis* | Fukiagehama Beach, Kagoshima, Japan |
| 84 | *Anadara inaequivalvis* | Fukiagehama Beach, Kagoshima, Japan |
| 85 | *Anadara inaequivalvis* | Fukiagehama Beach, Kagoshima, Japan |
| 86 | *Anadara inaequivalvis* | Fukiagehama Beach, Kagoshima, Japan |
| 87 | *Anadara inaequivalvis* | Fukiagehama Beach, Kagoshima, Japan |
| 88 | *Anadara inaequivalvis* | Fukiagehama Beach, Kagoshima, Japan |
| 89 | *Anadara inaequivalvis* | Fukiagehama Beach, Kagoshima, Japan |
| 90 | *Anadara inaequivalvis* | Fukiagehama Beach, Kagoshima, Japan |
| 91 | *Anadara inaequivalvis* | Fukiagehama Beach, Kagoshima, Japan |
| 92 | *Anadara inaequivalvis* | Fukiagehama Beach, Kagoshima, Japan |
| 93 | *Anadara inaequivalvis* | Fukiagehama Beach, Kagoshima, Japan |
| 94 | *Anadara inaequivalvis* | Fukiagehama Beach, Kagoshima, Japan |
| 95 | *Anadara inaequivalvis* | Fukiagehama Beach, Kagoshima, Japan |
| 96 | *Anadara inaequivalvis* | Fukiagehama Beach, Kagoshima, Japan |
| 97 | *Anadara inaequivalvis* | Fukiagehama Beach, Kagoshima, Japan |
| 98 | *Anadara inaequivalvis* | Fukiagehama Beach, Kagoshima, Japan |
| 99 | *Anadara inaequivalvis* | Fukiagehama Beach, Kagoshima, Japan |
| 100 | *Anadara inaequivalvis* | Fukiagehama Beach, Kagoshima, Japan |
| 101 | *Anadara inaequivalvis* | Fukiagehama Beach, Kagoshima, Japan |
| 102 | *Anadara inaequivalvis* | Fukiagehama Beach, Kagoshima, Japan |
| 103 | *Anadara inaequivalvis* | Fukiagehama Beach, Kagoshima, Japan |
| 104 | *Anadara inaequivalvis* | Fukiagehama Beach, Kagoshima, Japan |
| 105 | *Anadara inaequivalvis* | Fukiagehama Beach, Kagoshima, Japan |
| 106 | *Anadara inaequivalvis* | Fukiagehama Beach, Kagoshima, Japan |
| 107 | *Anadara inaequivalvis* | Fukiagehama Beach, Kagoshima, Japan |
| 108 | *Anadara inaequivalvis* | Fukiagehama Beach, Kagoshima, Japan |

STable. I: Complete list of specimens examined in this study, grouped by species.

| No. | Scientific Name | Locality |
|---|---|---|
| 109 | *Anadara inaequivalvis* | Fukiagehama Beach, Kagoshima, Japan |
| 110 | *Anadara inaequivalvis* | Fukiagehama Beach, Kagoshima, Japan |
| 111 | *Anadara inaequivalvis* | Fukiagehama Beach, Kagoshima, Japan |
| 112 | *Anadara inaequivalvis* | Fukiagehama Beach, Kagoshima, Japan |
| 113 | *Anadara inaequivalvis* | Fukiagehama Beach, Kagoshima, Japan |
| 114 | *Anadara inaequivalvis* | Fukiagehama Beach, Kagoshima, Japan |
| 115 | *Anadara inaequivalvis* | Fukiagehama Beach, Kagoshima, Japan |
| 116 | *Anadara inaequivalvis* | Fukiagehama Beach, Kagoshima, Japan |
| 117 | *Anadara inaequivalvis* | Fukiagehama Beach, Kagoshima, Japan |
| 118 | *Anadara inaequivalvis* | Fukiagehama Beach, Kagoshima, Japan |
| 119 | *Anadara inaequivalvis* | Fukiagehama Beach, Kagoshima, Japan |
| 120 | *Anadara inaequivalvis* | Fukiagehama Beach, Kagoshima, Japan |
| 121 | *Anadara inaequivalvis* | Fukiagehama Beach, Kagoshima, Japan |
| 122 | *Anadara inaequivalvis* | Fukiagehama Beach, Kagoshima, Japan |
| 123 | *Anadara inaequivalvis* | Fukiagehama Beach, Kagoshima, Japan |
| 124 | *Anadara inaequivalvis* | Fukiagehama Beach, Kagoshima, Japan |
| 125 | *Anadara inaequivalvis* | Fukiagehama Beach, Kagoshima, Japan |
| 126 | *Anadara inaequivalvis* | Fukiagehama Beach, Kagoshima, Japan |
| 127 | *Anadara inaequivalvis* | Fukiagehama Beach, Kagoshima, Japan |
| 128 | *Anadara inaequivalvis* | Fukiagehama Beach, Kagoshima, Japan |
| 129 | *Anadara inaequivalvis* | Fukiagehama Beach, Kagoshima, Japan |
| 130 | *Anadara inaequivalvis* | Fukiagehama Beach, Kagoshima, Japan |
| 131 | *Anadara inaequivalvis* | Fukiagehama Beach, Kagoshima, Japan |
| 132 | *Anadara inaequivalvis* | Fukiagehama Beach, Kagoshima, Japan |
| 133 | *Anadara inaequivalvis* | Fukiagehama Beach, Kagoshima, Japan |
| 134 | *Anadara inaequivalvis* | Fukiagehama Beach, Kagoshima, Japan |
| 135 | *Anadara inaequivalvis* | Fukiagehama Beach, Kagoshima, Japan |

STable. I: Complete list of specimens examined in this study, grouped by species.

| No. | Scientific Name | Locality |
|---|---|---|
| 136 | *Anadara inaequivalvis* | Fukiagehama Beach, Kagoshima, Japan |
| 137 | *Anadara inaequivalvis* | Fukiagehama Beach, Kagoshima, Japan |
| 138 | *Anadara inaequivalvis* | Fukiagehama Beach, Kagoshima, Japan |
| 139 | *Anadara inaequivalvis* | Fukiagehama Beach, Kagoshima, Japan |
| 140 | *Anadara inaequivalvis* | Fukiagehama Beach, Kagoshima, Japan |
| 141 | *Anadara inaequivalvis* | Fukiagehama Beach, Kagoshima, Japan |
| 142 | *Anadara inaequivalvis* | Fukiagehama Beach, Kagoshima, Japan |
| 143 | *Anadara inaequivalvis* | Fukiagehama Beach, Kagoshima, Japan |
| 144 | *Anadara inaequivalvis* | Fukiagehama Beach, Kagoshima, Japan |
| 145 | *Anadara inaequivalvis* | Fukiagehama Beach, Kagoshima, Japan |
| 146 | *Anadara inaequivalvis* | Fukiagehama Beach, Kagoshima, Japan |
| 147 | *Anadara inaequivalvis* | Fukiagehama Beach, Kagoshima, Japan |
| 148 | *Anadara inaequivalvis* | Fukiagehama Beach, Kagoshima, Japan |
| 149 | *Anadara inaequivalvis* | Fukiagehama Beach, Kagoshima, Japan |
| 150 | *Anadara inaequivalvis* | Fukiagehama Beach, Kagoshima, Japan |
| 151 | *Anadara inaequivalvis* | Fukiagehama Beach, Kagoshima, Japan |
| 152 | *Anadara inaequivalvis* | Fukiagehama Beach, Kagoshima, Japan |
| 153 | *Anadara inaequivalvis* | Fukiagehama Beach, Kagoshima, Japan |
| 154 | *Anadara inaequivalvis* | Fukiagehama Beach, Kagoshima, Japan |
| 155 | *Anadara inaequivalvis* | Fukiagehama Beach, Kagoshima, Japan |
| 156 | *Anadara inaequivalvis* | Fukiagehama Beach, Kagoshima, Japan |
| 157 | *Anadara inaequivalvis* | Fukiagehama Beach, Kagoshima, Japan |
| 158 | *Anadara inaequivalvis* | Fukiagehama Beach, Kagoshima, Japan |
| 159 | *Anadara inaequivalvis* | Fukiagehama Beach, Kagoshima, Japan |
| 160 | *Anadara inaequivalvis* | Fukiagehama Beach, Kagoshima, Japan |
| 161 | *Anadara inaequivalvis* | Fukiagehama Beach, Kagoshima, Japan |
| 162 | *Anadara inaequivalvis* | Fukiagehama Beach, Kagoshima, Japan |

STable. I: Complete list of specimens examined in this study, grouped by species.

| No. | Scientific Name | Locality |
|---|---|---|
| 163 | *Anadara inaequivalvis* | Fukiagehama Beach, Kagoshima, Japan |
| 164 | *Anadara inaequivalvis* | Fukiagehama Beach, Kagoshima, Japan |
| 165 | *Anadara inaequivalvis* | Fukiagehama Beach, Kagoshima, Japan |
| 166 | *Anadara inaequivalvis* | Fukiagehama Beach, Kagoshima, Japan |
| 167 | *Anadara inaequivalvis* | Fukiagehama Beach, Kagoshima, Japan |
| 168 | *Anadara inaequivalvis* | Fukiagehama Beach, Kagoshima, Japan |
| 169 | *Anadara inaequivalvis* | Fukiagehama Beach, Kagoshima, Japan |
| 170 | *Anadara inaequivalvis* | Fukiagehama Beach, Kagoshima, Japan |
| 171 | *Anadara inaequivalvis* | Fukiagehama Beach, Kagoshima, Japan |
| 172 | *Anadara inaequivalvis* | Fukiagehama Beach, Kagoshima, Japan |
| 173 | *Anadara inaequivalvis* | Fukiagehama Beach, Kagoshima, Japan |
| 174 | *Anadara kagoshimensis* | Katase-Higashi Beach, Kanagawa, Japan |
| 175 | *Anadara kagoshimensis* | Katase-Higashi Beach, Kanagawa, Japan |
| 176 | *Anadara kagoshimensis* | Ariake Sea, Kumamoto, Japan |
| 177 | *Anadara kagoshimensis* | Ariake Sea, Kumamoto, Japan |
| 178 | *Anadara kagoshimensis* | Ariake Sea, Kumamoto, Japan |
| 179 | *Anadara kagoshimensis* | Ariake Sea, Kumamoto, Japan |
| 180 | *Anadara kagoshimensis* | Aio-Higashi, Yamaguchi, Japan |
| 181 | *Anadara kagoshimensis* | Aio-Higashi, Yamaguchi, Japan |
| 182 | *Anadara kagoshimensis* | Aio-Higashi, Yamaguchi, Japan |
| 183 | *Anadara kagoshimensis* | Aio-Higashi, Yamaguchi, Japan |
| 184 | *Anadara kagoshimensis* | Aio-Higashi, Yamaguchi, Japan |
| 185 | *Anadara kagoshimensis* | Aio-Higashi, Yamaguchi, Japan |
| 186 | *Anadara kagoshimensis* | Aio-Higashi, Yamaguchi, Japan |
| 187 | *Anadara kagoshimensis* | Aio-Higashi, Yamaguchi, Japan |
| 188 | *Anadara kagoshimensis* | Aio-Higashi, Yamaguchi, Japan |
| 189 | *Anadara kagoshimensis* | Aio-Higashi, Yamaguchi, Japan |

STable. I: Complete list of specimens examined in this study, grouped by species.

| No. | Scientific Name | Locality |
| --- | --- | --- |
| 190 | *Anadara kagoshimensis* | Aio-Higashi, Yamaguchi, Japan |
| 191 | *Anadara kagoshimensis* | Aio-Higashi, Yamaguchi, Japan |
| 192 | *Anadara kagoshimensis* | Aio-Higashi, Yamaguchi, Japan |
| 193 | *Anadara kagoshimensis* | Aio-Higashi, Yamaguchi, Japan |
| 194 | *Anadara kagoshimensis* | Aio-Higashi, Yamaguchi, Japan |
| 195 | *Anadara kagoshimensis* | Mihama, Aichi, Japan |
| 196 | *Anadara kagoshimensis* | Mihama, Aichi, Japan |
| 197 | *Anadara kagoshimensis* | Mihama, Aichi, Japan |
| 198 | *Anadara kagoshimensis* | Mihama, Aichi, Japan |
| 199 | *Anadara kagoshimensis* | Nagasu, Kumamoto, Japan |
| 200 | *Anadara kagoshimensis* | Nagasu, Kumamoto, Japan |
| 201 | *Anadara kagoshimensis* | Surigahama, Ibusuki, Kagoshima, Japan |
| 202 | *Anadara kagoshimensis* | Surigahama, Ibusuki, Kagoshima, Japan |
| 203 | *Anadara kagoshimensis* | Unknown |
| 204 | *Anadara kagoshimensis* | Unknown |
| 205 | *Anadara kagoshimensis* | Unknown |
| 206 | *Anadara kagoshimensis* | Unknown |
| 207 | *Anadara kagoshimensis* | Unknown |
| 208 | *Anadara kagoshimensis* | Unknown |
| 209 | *Anadara satowi* | Zushi Beach, Kanagawa, Japan |
| 210 | *Anadara satowi* | Zushi Beach, Kanagawa, Japan |
| 211 | *Anadara satowi* | Zushi Beach, Kanagawa, Japan |
| 212 | *Anadara satowi* | Zushi Beach, Kanagawa, Japan |
| 213 | *Anadara satowi* | Zushi Beach, Kanagawa, Japan |
| 214 | *Anadara satowi* | Zushi Beach, Kanagawa, Japan |
| 215 | *Anadara satowi* | Zushi Beach, Kanagawa, Japan |
| 216 | *Anadara satowi* | Zushi Beach, Kanagawa, Japan |

STable. I: Complete list of specimens examined in this study, grouped by species.

| No. | Scientific Name | Locality |
|---|---|---|
| 217 | *Anadara satowi* | Zushi Beach, Kanagawa, Japan |
| 218 | *Anadara satowi* | Zushi Beach, Kanagawa, Japan |
| 219 | *Anadara satowi* | Zushi Beach, Kanagawa, Japan |
| 220 | *Anadara satowi* | Zaimokuza Beach, Kanagawa, Japan |
| 221 | *Anadara satowi* | Zaimokuza Beach, Kanagawa, Japan |
| 222 | *Anadara satowi* | Katase-Higashi Beach, Kanagawa, Japan |
| 223 | *Anadara satowi* | Katase-Higashi Beach, Kanagawa, Japan |
| 224 | *Anadara satowi* | Katase-Higashi Beach, Kanagawa, Japan |
| 225 | *Anadara satowi* | Fukiagehama Beach, Kagoshima, Japan |
| 226 | *Anadara satowi* | Tsuyazaki, Fukuoka, Japan |
| 227 | *Anadara satowi* | Kujukuri Beach, Chiba, Japan |
| 228 | *Anadara satowi* | Kujukuri Beach, Chiba, Japan |
| 229 | *Anadara satowi* | Kujukuri Beach, Chiba, Japan |
| 230 | *Anadara satowi* | Kujukuri Beach, Chiba, Japan |
